# Taxonomic propagation of phenotypic features predict host pathogen interactions

**DOI:** 10.1101/508762

**Authors:** Wang Liu-Wei, Şenay Kafkas, Robert Hoehndorf

**Affiliations:** Computer, Electrical and Mathematical Sciences and Engineering Division, Computational Bioscience Research Center, King Abdullah University of Science and Technology, Thuwal 23955, Saudi Arabia

## Abstract

Identification of host-pathogen interactions can reveal mechanistic insights of infectious diseases for potential treatments and drug discoveries. Current computational methods focus on the prediction of host–pathogen protein interactions and rely on our knowledge of the sequences and functions of pathogen proteins, which is limited for many species, especially for emerging pathogens. We developed an ontology-based machine learning method that predicts potential interaction protein partners for pathogen taxa. Our method exploits information about infectious disease mechanisms through features learned from phenotypic, functional and taxonomic knowledge about pathogen taxa and human proteins. Additionally, by propagating the phenotypic features of the pathogens within a formal representation of pathogen taxonomy, we demonstrate that our model can also accurately predict interaction protein partners for pathogens even without known phenotypes, using a combination of their taxonomic relationships with other pathogens and information from ontologies as background knowledge. Our results show that the integration of phenotypic, functional and taxonomic knowledge not only improves the prediction performance, but also enables us to investigate novel pathogens in emerging infectious diseases.

**Author summary:** Infectious diseases are caused by various types of pathogens, such as bacteria and viruses, and lead to millions of deaths each year, especially in low-income countries. Researchers have been attempting to predict and study possible host-pathogen interactions on a molecular level. Understanding these interactions can shed light on how pathogens invade cells or disrupt the immune system. We propose a novel method to predict such interactions by associating phenotypes (e.g., the signs and symptoms of patients) associated with pathogens and phenotypes associated with human proteins. We are able to accurately predict and prioritize possible protein partners for dozens of pathogens. We further extended the prediction model by relating pathogens without phenotypes with those with phenotypes through their taxonomic relationships. We found that the addition of taxonomic knowledge greatly increased the number of pathogens that we can study, without diminishing the accuracy of the model. To the best of our knowledge, we are the first to predict host-pathogen interactions based on phenotypes and taxonomy. Our work has important implications for new pathogens and emerging infectious diseases that are not yet well-studied.

## Introduction

Infectious diseases from bacteria, viruses and fungi are one of the major causes of deaths around the globe, especially in low-income regions [1]. Pathogens disrupt host cell functions [2] and target immune pathways [3] through complex interactions among proteins [4], RNA [5] and DNA [6]. The study of host-pathogen interactions (HPI) can provide insights into molecular mechanisms underlying infectious diseases and enable the development of novel therapeutic discoveries. For example, a previous study of many HPIs showed that pathogens typically interact with the protein hubs (those with many interaction partners) and bottlenecks (those of central location to important pathways) in human protein-protein interaction (PPI) networks [4]. However, due to high costs and time constraints, experimentally validated pairs of interacting host-pathogen proteins are limited in number. Moreover, there exists a time delay for a validated HPI to be included in a database of HPIs, often requiring text-mining efforts [7]. Therefore, the computational prediction of HPIs is useful in suggesting candidate interaction partners out of all the human proteins.

Existing HPI prediction methods typically focus on predicting protein interactions and utilize features of interacting protein pairs, such as PPI network topology, structural and sequential homology, or functional profiling such as Gene Ontology similarity and KEGG pathway analysis [8]. While such protein-specific features exist for some extensively studied pathogens, such as *Mycobacterium tuberculosis* [9], human immunodeficiency virus [10], *Salmonella* and *Yersinia* [11], for most of the other pathogens, especially the newly emerging ones, these features are scarce (or non-existent) and expensive to obtain. As new virus species are discovered each year with potentially many more to come [12], a method is needed to aggregate our existing phenotypic, functional and taxonomic knowledge about HPIs and predict for pathogens that are less well studied or even novel.

The phenotypes elicited by pathogens, i.e., the signs and symptoms observed in a patient, may provide information about molecular mechanisms [13]. The information that phenotypes provide about molecular mechanisms is commonly exploited in computational studies of Mendelian disease mechanisms [14, 15], for example to suggest candidate genes [16, 17] or diagnose patients [18], but can also be used to identify drug targets [19] or gene functions [20]. To the best of our knowledge, phenotypes and phenotype similarity have not yet been used to suggest host-pathogen interactions.

We use the hypothesis that phenotypes elicited by an infection with a pathogen are, among others, the result of molecular interactions, and that knowledge of the phenotypes in the host can be used to suggest the protein perturbations through which these phenotypes arise. While a large number of phenotypes resulting from infections are a consequence of immune system processes that are shared across a wide range of different types of pathogens, certain hallmark phenotypes, such as decreased CD4 count in infections with human immunodeficiency virus or microcephaly resulting from Zika virus infections, can be used to suggest candidate host proteins through which these phenotypes are elicited. Phenotypes shared across many pathogens could also contribute to the prediction of interacting host proteins as well as the absence of certain phenotypes.

We developed a method to predict candidate interacting proteins for pathogens by integrating phenotypic, functional and taxonomic information about pathogens and human proteins. Relying on recent progress in deep learning with structured and semantic data [21], we construct a model that combines this information as well as structured information about taxonomic relations between pathogens to predict host-pathogen interactions. We demonstrate that our model can not only accurately predict the interacting protein partners for pathogens with phenotypic information, but also their relatives without phenotypes by aggregating knowledge through the taxonomy.

## Results

### Feature generation and representation learning for human proteins and pathogens

We first generated feature vectors for pathogens and human proteins based on different sets of features, and then trained an artificial neural network model (ANN) for predicting their interactions. We follow the workflow shown in Figure 1.

**Fig 1.**
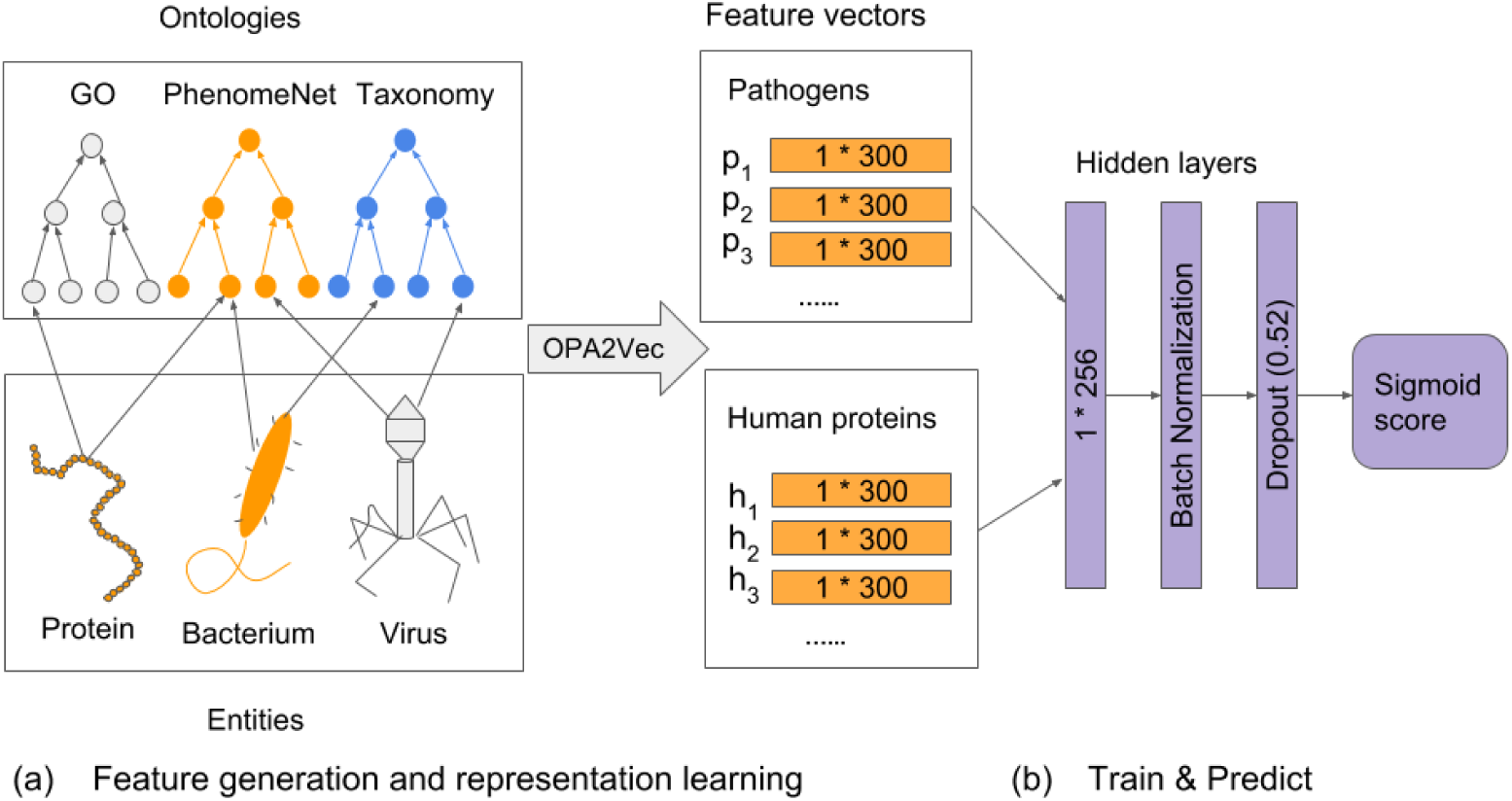
An illustration of our overall workflow.

The first step in our method is the generation of features that represent a pathogen or a human protein. We represent human proteins through their associated phenotypes, the phenotypes associated with their mouse orthologs, and their functions. We obtained loss-of-function phenotypes of mouse genes from the Mouse Genome Informatics (MGI) database [22] and phenotype annotations of human genes from Human Phenotype Ontology (HPO) database [23]. We obtained protein function annotations from the Gene Ontology (GO) database [24, 25]. For pathogens, we use the phenotype annotations from the PathoPhenoDB [26], a database of pathogen-phenotype associations, and taxonomic information from the NCBI Taxonomy [27].

We represent the phenotypes, functions and taxonomic relations through ontologies. To associate human and mouse phenotypes, we use the cross-species phenotype ontology PhenomeNET [16, 28], which combines the Human Phenotype Ontology (HP) [23] and the Mammalian Phenotype Ontology (MP) [29]. To incorporate knowledge of protein functions, we use the Gene Ontology [24, 25]. For the taxonomic relations between pathogens, we used the NCBI Taxonomy Ontology [27]. These ontologies contain formalized biological background knowledge [30]. Using the information in ontologies as background knowledge during feature generation has the potential to significantly improve the performance of these features in machine learning and predictive analyses [31].

We use the OPA2Vec [21] method to generate features from annotations of pathogen taxa and human proteins while using the ontologies that are used to express them as background knowledge. To determine the effect of different annotations and ontologies on the performance of our models, we generated feature vectors of human proteins from human phenotype annotations, the phenotypes of their mouse orthologs, and functional annotations individually with their respective ontologies, as well as their union and intersection by combining the annotations and merging the ontologies.

### Phenotypic, functional and taxonomic prediction of interaction partners

We first test our method on pathogens that have phenotype annotations in PathoPhenoDB and use a merged representation of the PhenomeNet and NCBI Taxonomy ontologies. We obtain high-confidence interaction pairs of pathogens and human proteins from the Host-Pathogen Interaction Database (HPIDB) [32] as the positive training data. We then train an ANN to predict host-pathogen interactions using the feature vectors of pathogens and human proteins as input. We perform leave-one-taxon-out cross-validation (LOOCV), in which we leave out one pathogen taxon in our positive set for validation, and use all the other pathogens and their interactions as training data. We evaluated our model by ranking the true positive protein partners of the validation taxon (see Methods). We perform the training and evaluation separately based on taxonomic groups, i.e., viruses and bacteria. Table 1 provides an overview over the underlying data used for these predictions, and Figure 2 shows the prediction results.

**Table 1.** Datasets used for phenotypic, functional and taxonomic predictions. The features of human proteins are human phenotypes (HP), mouse phenotypes (MP), gene ontology terms (GO), their union and their intersection. The features of pathogens are phenotypes from PathoPhenoDB and taxonomic relations from NCBI Taxonomy. The coverage shows the total number of proteins annotated with the respective features. The number of unique interacting pairs, pathogen taxa and human proteins are listed for both viruses and bacteria.

| Features | Coverage | Viruses |  |  | Bacteria |  |  |
| --- | --- | --- | --- | --- | --- | --- | --- |
|  |  | Interactions | Taxa | Proteins | Interactions | Taxa | Proteins |
| HP | 3740 | 101 | 24 | 78 | 18 | 9 | 18 |
| MP | 9153 | 214 | 29 | 174 | 47 | 15 | 46 |
| GO | 18084 | 291 | 30 | 247 | 57 | 16 | 56 |
| HP∪MP∪GO | 18349 | 291 | 30 | 247 | 57 | 16 | 56 |
| HP∩MP∩GO | 2955 | 92 | 23 | 69 | 18 | 9 | 18 |

**Fig 2.**
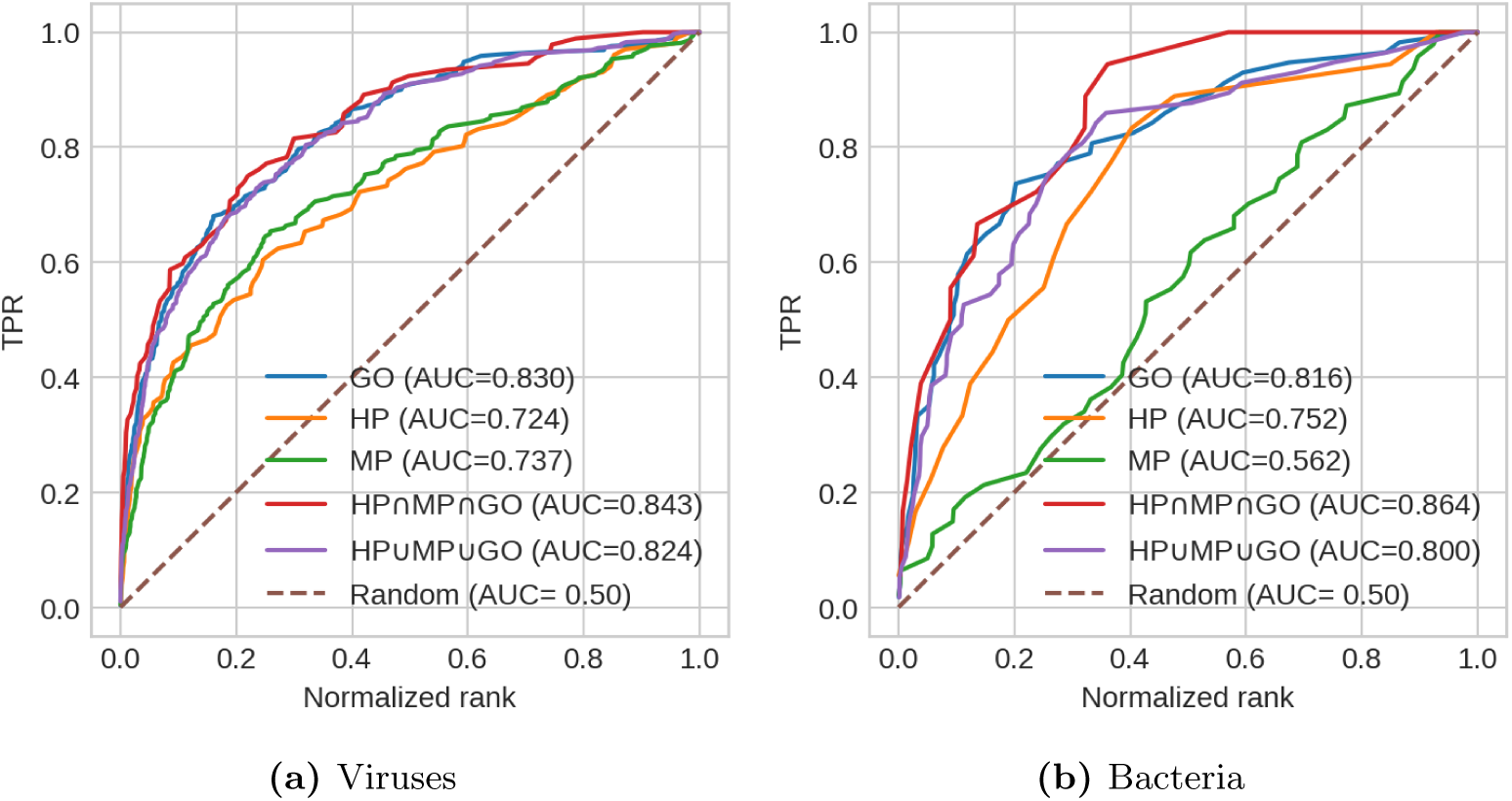
The TPR curves by normalized rank of proteins. The solid curves represent the different features used to generate the feature vectors of human proteins. The dashed line represents a random classifier.

For both viruses and bacteria, the model using the intersection of phenotypic and functional features has the best performance in prioritizing true protein partners. This model is able to prioritize 50% of the true positives within the first 10% of all proteins. The TPR curve of bacteria is less smooth compared to the viruses due to the limited number of exact matches of bacterial taxa between HPIDB and PathoPhenoDB. To compare with the effect of only using phenotypic features, we also generated feature vectors for pathogens only using phenotypes and included the performance in S1 File.

### Propagation of phenotypic features via pathogen taxonomy

The previous section only considers pathogens that have phenotypic annotations from the associated infectious diseases in PathoPhenoDB, which presents a problem of missing data: many pathogen taxa in HPIDB cannot be matched with those in PathoPhenoDB at the exact taxon level and are ignored during training. To address the taxonomic inconsistencies, we propagate information across a taxonomy: since we use a merged representation from the PhenomeNet and NCBI Taxonomy ontologies during feature generation, there can always be a representation of phenotypes for any pathogen taxon within the taxonomic hierarchy. The representation learned from taxonomy not only increases the taxonomic coverage but also encodes phenotypic and taxonomic information for pathogens and their relatives by exploiting the underlying ontological structures. Table 2 shows that almost all of the pathogens we consider have some relatives within the same family that have some phenotypic annotations in PathoPhenoDB, and over two thirds within the same species (i.e., species, subspecies, or strain).

**Table 2.** Coverage of phenotypic features through taxonomy. The percentages stand for the proportion of pathogens that have relatives within the same family, genus or species with phenotypic annotations and thus, have a vector representation.

| Features | Viruses |  |  | Bacteria |  |  |
| --- | --- | --- | --- | --- | --- | --- |
|  | Family | Genus | Species | Family | Genus | Species |
| HP | 100% | 90% | 77% | 97% | 97% | 79% |
| MP | 100% | 90% | 76% | 98% | 98% | 83% |
| GO & HP∪MP∪GO | 100% | 89% | 75% | 98% | 98% | 83% |
| HP∩MP∩GO | 100% | 90% | 75% | 97% | 97% | 79% |

While we want to learn phenotype representations from taxonomy, the training data obtained from HPIDB may have pathogens that are closely related taxa; therefore, the taxonomic propagation could simply transfer the information and make the prediction task trivial and unrealistic to be generalizable for novel pathogens. To simulate what we would expect to see for a novel pathogen that does not have any well-studied close relatives, we further excluded from the training data, at each taxon fold, the interactions from the same viral family for viruses and the same bacterial genus for bacteria.

We retrained the models with the extended set of pathogens and evaluated their performance using LOOCV. Table 3 shows that there is more than 50% increase in the number of interactions and pathogen taxa after the taxonomic propagation. Figure 3 shows the model performance after retraining with the extended set of pathogens. For bacteria, the prediction model shows an increase in AUC for every feature of human proteins and for viruses the performance remains on a similar level. Again, the intersection of phenotypic and functional features outperforms the others while the GO features performs better than other individual features. Table 4 shows a summary of the performances with and without the taxonomic propagation. We also test the model performance without excluding close relatives and show the performance in S1 File.

**Table 3.**
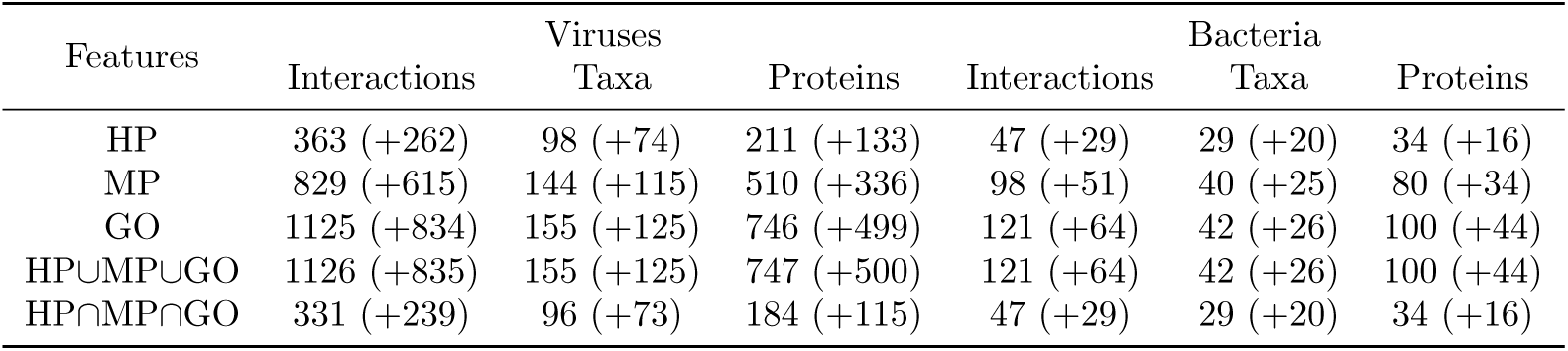
Datasets used for predictions via taxonomy. The numbers in brackets indicate the increase in data after using the propagated representations via pathogen taxonomy.

**Table 4.** A summary of the model performances. The bold AUC values represent, for each feature of human proteins and for virus and bacteria, the best model performance before or after using the propagated representations via taxonomy.

| Features | Without propogation |  | With propagation |  |
| --- | --- | --- | --- | --- |
|  | Virus | Bacteria | Virus | Bacteria |
| HP | **0.724** | 0.752 | 0.722 | **0.814** |
| MP | **0.737** | 0.562 | 0.717 | **0.683** |
| GO | 0.830 | 0.816 | **0.837** | **0.883** |
| HP∪MP∪GO | 0.824 | 0.800 | **0.825** | **0.862** |
| HP∩MP∩GO | **0.843** | 0.864 | 0.841 | **0.908** |

**Fig 3.**
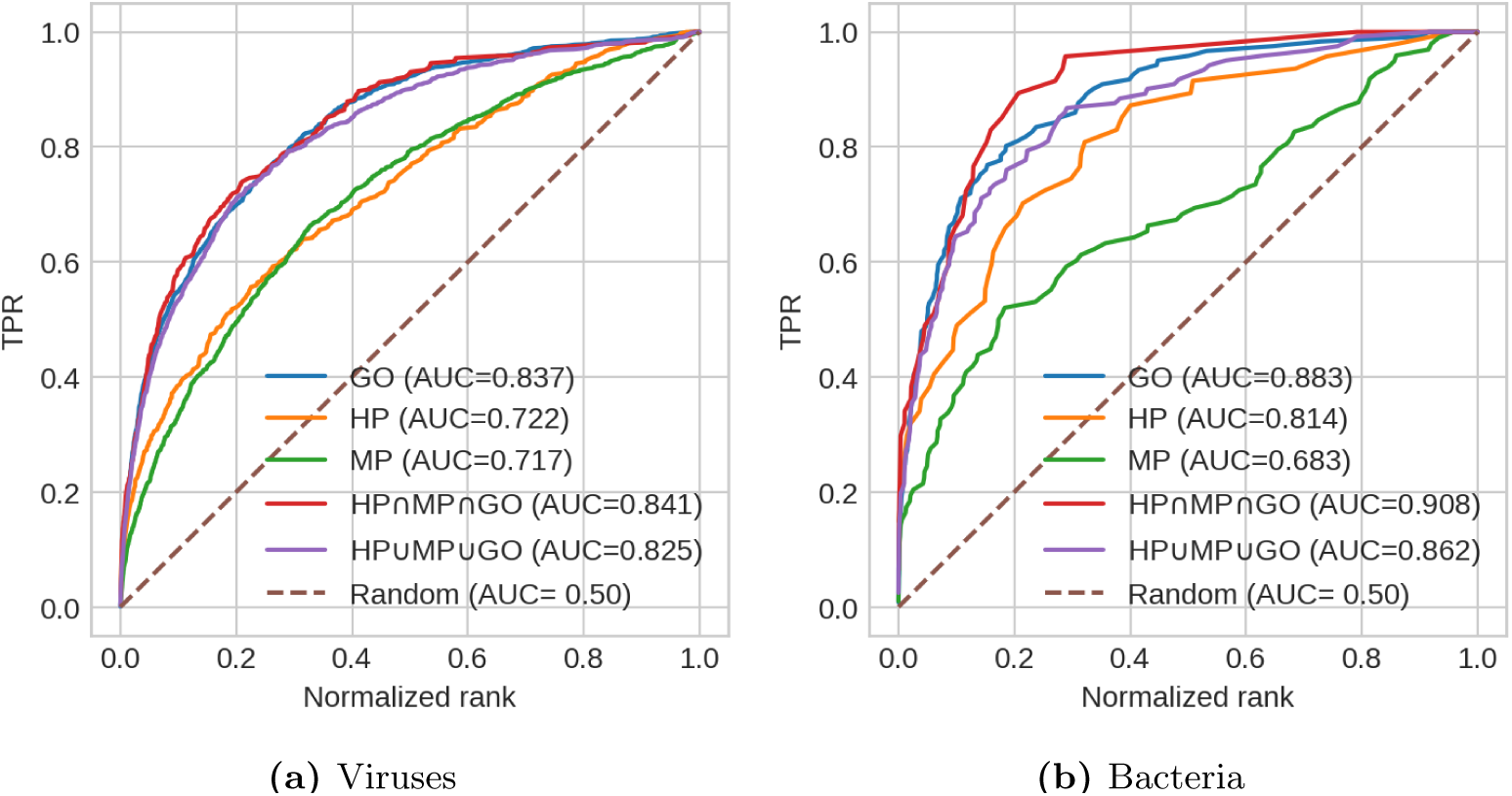
The TPR curves by normalized rank of proteins after using the propagated representations of phenotypes via the taxonomy of pathogens.

## Discussion

We developed an ontology-based method for predicting HPIs using the phenotypic, functional and taxonomic features of pathogens and human proteins. Unlike previous methods that target protein-protein interactions and utilize features from protein structures and sequences [8], we focused on predicting pathogen-protein interactions and use features that are relatively easy and fast to obtain from patients’ symptoms and taxonomic identification of the pathogens.

Since the existing HPI prediction methods commonly predict interactions between proteins (protein-protein interactions), our model of pathogen-protein interactions is not directly comparable. However, we compare our results with the protein-based prediction methods by combining the predicted human proteins for each pathogen across several studies [33–35] and ranking them with our predicted score. The predicted protein partners by these methods [33–35] are ranked significantly higher than other proteins (Mann-Whitney U test; HIV: *p* = 6.57 × 10^−23^; HBV: *p* = 9.65 × 10^−4^; *B. anthracis*: *p* = 0.014; *F. tularensis*: *p* = 5.68 × 10^−7^; *Y. pestis*: *p* = 0.02; *S. typhi*: *p* = 4.19 × 10^−8^). This suggests that, although we are using very different features, our method captures some common molecular mechanisms underlying pathogenesis that are also predicted by protein-based prediction methods.

Phenotype-based approaches have been successful in predicting disease-gene associations for Mendelian diseases [16] but have not yet been used for prediction of host-pathogen interactions for infectious diseases. Our results demonstrate that the phenotypes associated with human proteins and protein functions (i.e., GO classes), can generate accurate predictions. While GO classes have been shown to perform well in HPI predictions [36, 37], they still rely on the sequence homology of pathogen proteins. We only target interactions between pathogen taxa and host proteins instead of direct protein-protein interactions, and therefore avoid the common bottleneck of identifying the functions of pathogen proteins. Instead, we introduce a novel and – in the context of infectious diseases – rarely explored type of feature, the phenotypes elicited by pathogens in their hosts, as a “proxy” for the molecular mechanisms that result in the phenotypes.

The focus of our method on utilizing features generated based on endo-phenotypes observed in humans and mice [38] has the crucial advantage that we can identify host-pathogen interactions that may contribute to particular signs and symptoms. For example, our model predicts an interaction between Zika virus (`NCBITaxon:64320`) and DDX3X (`UniProt:O00571`). Infections with Zika virus have the potential to result in abnormal embryogenesis and, specifically, microcephaly [39]. Phenotypes associated with DDX3X in the mouse ortholog include abnormal embryogenesis, microcephaly, and abnormal neural tube closure [40]. DDX3X mutations in humans have been found to result in intellectual disability, specifically in females and affecting individuals in dose-dependent manner [41]. While DDX3X has previously been linked to the infectivity of Zika virus [42], our model further suggests a role of DDX3X in the development of the embryogenesis phenotypes resulting from Zika virus infections.

The lack of phenotypic annotations is often a challenge for such phenotype-based predictions, which is also the case for many pathogens in our study. We demonstrated that by integrating a formalized taxonomy in our feature generation and propagating phenotypic features via the pathogen taxonomy, not only did we greatly increase the coverage of pathogen taxa but also improved the prediction performance. This suggests that our model was able to learn high-level representations of the phenotypes caused by pathogens across their taxonomic hierarchy. This is the first attempt, to our knowledge, for a taxonomy ontology to be used in a machine learning method to transfer and distribute information across different taxonomic levels. This method has the potential to be extended to many applications where data are only present for some levels of taxonomy, such as the modeling of some other ecological systems where many species interact.

## Materials and methods

### Data sources

We downloaded interactions between hosts and pathogens from the Host Pathogen Interaction Database (HPIDB) [32] on July 16, 2018. The dataset contains 27,533 distinct interaction pairs of human proteins to pathogen species with the IntAct and Virhostnet MI scores provided. We further filtered the interactions by the MI scores with a threshold of 0.5, reducing the number of interaction pairs to 1,682. We obtained phenotypes associated with pathogens from the PathoPhenoDB [26], a database of manually curated and text-mined associations of pathogens, diseases and phenotypes. We downloaded the version 1 of the PathoPhenoDB database (http://patho.phenomebrowser.net/) on Oct 11, 2018. Similarly to the implementation at the PathoPhenoDB web interface, species level taxa are also annotated with the phenotypes of all their subclasses in the taxonomy of pathogens.

We downloaded phenotypes of human genes on August 26, 2018 from the Human Phenotype Ontology (HPO) database. The dataset contains 3,777 human genes and 131,045 associations between human genes and HPO classes. We downloaded phenotypes associated with mouse genes from the Mouse Genome Informatics (MGI) database [22] on September 17, 2018, using the `MGI_GenePheno.rpt` file with 12,182 mouse genes and 165,892 associations of mouse genes and Mammalian Phenotype ontology (MP) classes. We then associated MP classes with human genes by mapping mouse genes to human genes based on the information about orthologous genes provided in `HMD_HumanPhenotype.rpt` downloaded from MGI on July 16, 2018. We remove all “no abnormal phenotype detected” (`MP:0002169`) annotations.

We mapped all the genes to Uniprot proteins using the Uniprot Retrieve/ID mapping tool (https://www.uniprot.org/uploadlists), and used only reviewed Swissprot mappings. We downloaded Gene Ontology (GO) annotations of human proteins from the Gene Ontology website [24, 25] on September 27, 2018. To identify broad taxonomic groups, i.e., bacteria and viruses, we used the NCBI Taxonomy Categories file (ftp://ftp.ncbi.nih.gov/pub/taxonomy) downloaded on September 3, 2018.

To obtain formal representations of phenotypes and GO classes, we downloaded the cross-species PhenomeNET Ontology [16, 28] from the AberOWL ontology repository [43] on September 13, 2018, and we downloaded the Gene Ontology [24, 25] from the Gene Ontology website on September 27, 2018. For taxonomy features, we downloaded the NCBI Taxonomy in OWL format from the OBO Foundry website (http://www.obofoundry.org/ontology/ncbitaxon.html) on October 8th, 2018.

### Generation of feature embeddings

For feature generation, we used OPA2Vec [21], a tool that generates feature vectors from entities and their annotation from ontologies. OPA2vec first generates a corpus consisting of relevant annotations of pathogens and proteins (i.e., HP, MP, GO, and NCBI Taxonomy classes), the complete set of class axioms and the metadata of the ontologies. For OPA2Vec parameters, we used the default skipgram model with a window size of 5 and a minimum count parameter of 2. We tested different embedding sizes (i.e., 50, 100, 150, 200 and 300) and fixed the embedding size to 300 throughout our experiments in this paper.

### Supervised prediction models

We trained an ANN as a binary classifier that predicts a sigmoid score of an interaction between a host protein and a pathogen taxon, and we used leave-one-taxon-out cross-validation to report performance results. During each taxon-fold, we use the trained ANN to predict possible interacting proteins for the validation taxon. We evaluate the prediction results by ranking the sigmoid scores of its true protein partners among those of all the proteins. By aggregating the ranks of these true positive at each taxon fold, we plotted a true positive rate (TPR) curve by the normalized ranks of proteins and calculated an Area Under Curve (AUC) metric by trapezoidal rule for integration. The AUC metric shows our model’s ability to prioritize the true protein partners under the assumption that a perfectly random model will have an AUC of 0.5. Due to the large number of negatives in relation to the positives, normalized ranks approximate the true negative rate (TNR) and the AUC measure approximates the area under the receiver operating characteristic (ROC) curve [44].

We obtained positives by extracting interaction pairs of human proteins and pathogen taxa from HPIDB. To generate the negatives, we first generated all possible combinations between all the proteins with a feature vector and pathogens that are in the positives, and then randomly sampled within the ones that did not occur in the positives. We treat all “unknown” interactions as negatives.

We searched for an optimal set of hyperparameters using the Hyperas library (https://github.com/maxpumperla/hyperas) based on the TPE algorithm [45] for one of our models (i.e., the model using only the HP features). After searching, we fixed our parameters to the following throughout all the experiments: one hidden layer of 256 units with Rectified Linear Unit (ReLU) [46] as activation function; one batch normalization layer; one dropout layer with a dropout rate of 0.52; one output layer with a sigmoid activation function; the Adam optimizer with cross entropy as loss function. We provide the complete list of hyperparameter search space and plot the AUC by epoch relationship in S2 File. The preprocessing and training pipeline is provided as Jupyter Notebooks, which are available at https://github.com/bio-ontology-research-group/hpi-predict.

## Acknowledgements

We would like to thank Maxat Kulmanov and Mona Alshahrani for their advice on using machine learning libraries.

## Supporting information

**S1 File Model performance using only phenotypes for pathogens, and not excluding close relatives.**

**S2 File Hyperas search space and AUC-epoch plots.**

