## Supplementary material for "Taxonomic propagation of phenotypic features predict host pathogen interactions": S1 File

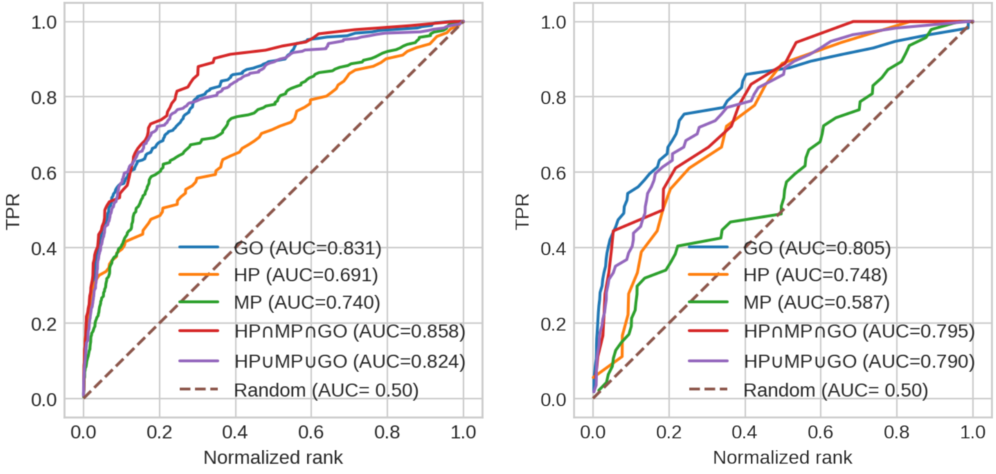


**(a)** Viruses **(b)** Bacteria

**Fig 1. The TPR curves by normalized rank of proteins only using phenotypes as features for pathogens**

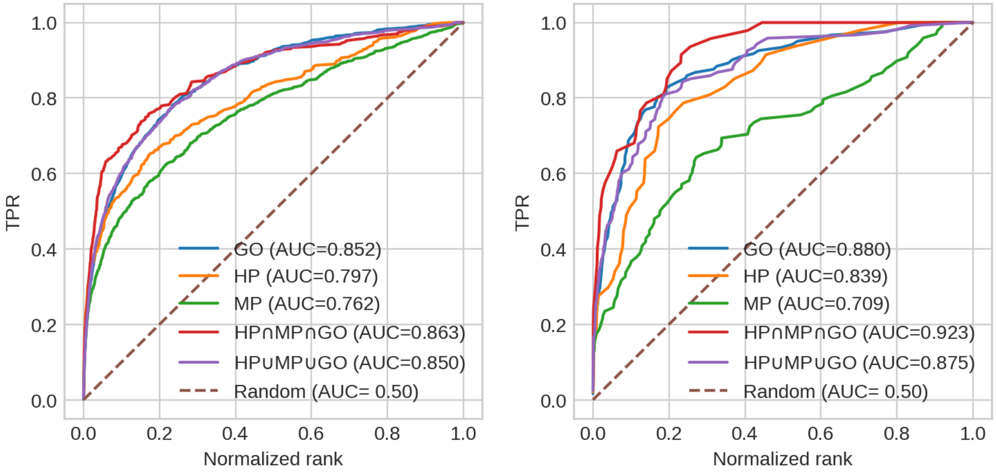


**(a)** Viruses **(b)** Bacteria

**Fig 2. The TPR curves by normalized rank of proteins without excluding close taxonomic relatives**
