## Supplementary material for "Taxonomic propagation of phenotypic features predict host pathogen interactions": S2 File

**Table 1. Hyperparameter search space for Hyperas Optimizer.**

| Dimension | First layer | Second layer | Third layer | Activation | Dropout | Optimizer |
|---|---|---|---|---|---|---|
| 50 | 64, 32, 16, 8, 4 | 16, 8, 4 | - | | | |
| 100 | 128, 64, 32, 16, 8 | 32, 16, 8 | 8, 4, 2 | | | |
| 150 | 256, 128, 64, 32, 16 | 32, 16, 8, 4 | 8, 4, 2 | Relu, Sigmoid | (0, 1) | Rmsprop, Adam |
| 200 | 256, 128, 64, 32, 16 | 32, 16, 8, 4 | 8, 4, 2 | | | |
| 300 | 512, 256, 128, 64, 32 | 64, 32, 16, 8 | 8, 4, 2 | | | |

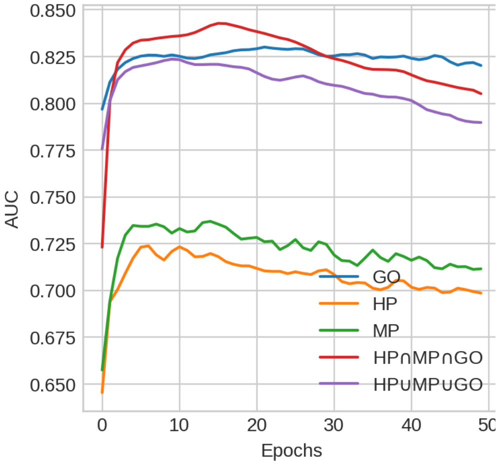


**(a)** Viruses (without propagation)

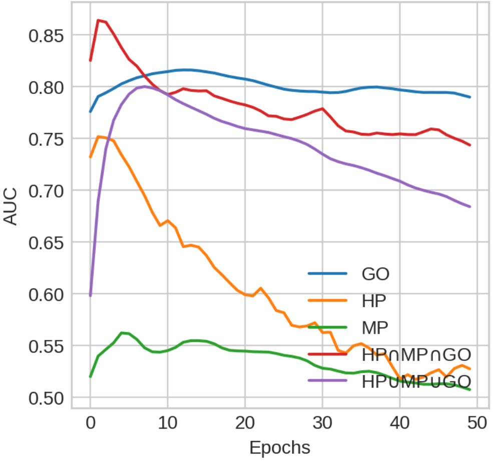


**(b)** Bacteria (without propagation)

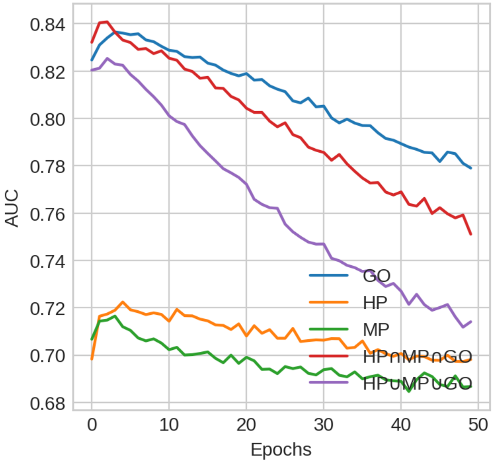


**(c)** Viruses (with propagation)

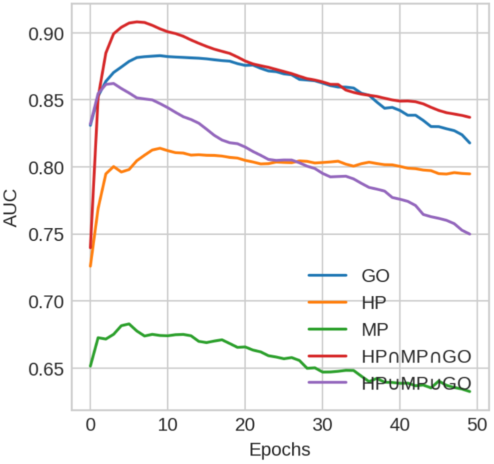


**(d)** Bacteria (with propagation)

**Fig 1. The AUC curves by the number of epochs of training**
